# The human-baited host decoy trap (HDT) is an efficient sampling device for exophagic malaria mosquitoes within irrigated lands in southern Malawi

**DOI:** 10.1101/2021.08.26.457772

**Authors:** Kennedy Zembere, James Chirombo, Peter Nasoni, Daniel P. McDermott, Lizzie Tchongwe-Divala, Frances M. Hawkes, Christopher M. Jones

## Abstract

Irrigation schemes provide an ideal habitat for *Anopheles* mosquitoes particularly during the dry season. Reliable estimates of outdoor host-seeking behaviour are needed to assess the impact of vector control options and this is particularly the case for *Anopheles arabiensis* which displays a wide range of behaviours that circumvent traditional indoor-insecticide based control. In this study we compared the Host Decoy Trap (HDT) with the Human Landing Catch (HLC) and Suna trap in a repeated Latin square design in two villages on an irrigated sugar estate in southern Malawi. Over the course of 18 trapping nights we caught 379 female *Anopheles*, the majority of which were identified as *An. arabiensis*. Overall, the HDT and HLC caught a similar number of *Anopheles* per night with both methods catching significantly higher densities than the Suna trap across both villages. Regardless of the density of *Anopheles* mosquitoes in each village the HLC and HDT demonstrated broadly similar sampling efficacy. We conclude that the HDT is an effective sampling device for outdoor host seeking *An. arabiensis* in southern Malawi. The presence of *An. arabiensis* in irrigated lands during the dry season poses a challenge for ongoing indoor vector control efforts.

## Introduction

Irrigation provides a refuge for malaria mosquito vectors (in the genus *Anopheles*) during periods of the year when environmental conditions would otherwise be too hot or too dry to sustain local mosquito populations. The flooding, seepage or spill over of water from irrigation and drainage channels creates the ideal microhabitat for the aquatic life stages of anophelines^1^. Depending on the agro-ecosystem, the cultivation of land can permanently shift local mosquito population dynamics and living in proximity to agricultural schemes is a risk factor for exposure to potentially infectious mosquito bites^2,3^

One of the main species to benefit from irrigated agriculture is *Anopheles arabiensis* (a member of the *Anopheles gambiae* species complex), particularly in eastern and southern parts of Africa. This species is generally associated with dry-savannah habitats and adults oviposit in temporary, small, sunlit pools^4^, meaning that irrigated cropland provides ideal conditions all year-round. In Kenya, for example, the pattern of larval and adult population densities of *An. arabiensis* reflect the rice cropping cycle with peak abundances occurring when rice stalks are immature after transplantation^5^. The shallow periphery of microdams – small water harvesting structures – support dominant *An. arabiensis* populations in Western Kenya^6^ and Ethiopia^7^, and several studies have observed increases in *An. arabiensis* density, entomological inoculation rates and malaria incidence as a function of distance from small or large agricultural schemes^7–9^.

Controlling *An. arabiensis* via contemporary vector control methods such as insecticide treated nets (ITNs) and indoor residual spraying (IRS) is compromised by the large amount of behavioural plasticity exhibited by this species^10^. *Anopheles arabiensis* displays a continuum of feeding (zoophagy and exophagy) and resting (exophily) patterns which can circumvent indoor insecticide-based control. In many areas of East Africa, including Malawi, *An. arabiensis* has superseded the more anthropophilic *An. gambiae s*.*s*. as the dominant vector of the *Anopheles gambiae* species complex following the upscale of ITNs^11^. Complementary interventions which target the full spectrum of *An. arabiensis* behaviour are needed to reduce residual transmission. This requires the simultaneous development of reliable and practical means of quantifying host-seeking behaviour to determine the efficacy of these interventions.

Unlike other crops, which are fallow for part of the year, sugarcane is a perennial crop which can be irrigated all year round^12^. This provides an ideal environment for malaria vectors to flourish and extend the transmission season beyond the typical window of the rainy season. Large commercial sugar estates process sugarcane at source, reducing transport and production costs. This encourages the movement of employees and families into the area increasing the local population density. The combination of year-round mosquito breeding and human settlements within and adjacent to the sugar fields creates a ‘*malaria microcosm*’ leading many private sugar-processing firms to invest in their own bespoke malaria control programmes as part of healthcare initiatives for the workforce^13^.

The Illovo Sugar company operates two estates in Malawi with the larger of these estates (Nchalo) located next to the Shire River in the southern district of Chikwawa. Malaria prevalence is historically high and the current *Plasmodium falciparum* prevalence in children aged 2-10 years old is estimated at 12.2%^14^. Illovo has conducted an IRS programme since 1990 to reduce malaria incidence^15^ and currently target all households with a single-application of the organophosphate, pirimiphos-methyl.

In this study, we quantify outdoor host-seeking malaria vectors in two villages situated inside the sugar estate boundary in Nchalo. We compare outdoor catches using three methods; the host decoy trap (HDT)^16^, human landing catch (HLC)^17^ and the Suna trap^18^. Our aim is to compare the performance of the recently developed HDT as an outdoor *Anopheles* sampling device in an area where the risk of exposure to *Anopheles* bites is perceptibly higher due to the irrigated habitat provided by the sugarcane fields. We measure outdoor host-seeking in the face of the high IRS coverage in Illovo and during the middle of the dry season as part of our efforts to understand the ecology of the main vectors in the area for improved control.

## Results

We collected a total of 2,096 mosquitoes over the course of 18 nights during the middle of the dry season (June-July 2019) on the Nchalo sugar estate using three outdoor sampling methods. Altogether, we identified 379 *Anopheles* and 1,717 culicines. We primarily focused our analyses on the 379 *Anopheles*, all of which were female.

There was a stark difference in the abundance of *Anopheles* caught between the two study villages despite their proximity (∼8 km apart) with a 90% reduction in Mwanza (RR = 0.10, P = 2e^-16^). Across both villages the HDT and HLC caught comparable numbers of *Anopheles* per night (RR = 0.85, P = 0.508) whereas significantly fewer *Anopheles* were caught by the Suna trap compared to the HLC (RR = 0.36, P = 0.0005). In Lengwe, the HDT sampled a nightly mean of 7.9 *Anopheles* (95% CI: 4.6-11.1) compared with 8.0 by HLC (95% CI: 4.3-11.8) while the Suna trap caught 2.8 *Anopheles* (Figure 2A). In Mwanza, where the overall number of *Anopheles* was considerably lower, the HDT caught a nightly average of 0.7 (95% CI: 0.002-1.44) compared with 1.2 by the HLC (95% CI: 0.037-0.63) whereas only 0.3 (CI: 0.056-2.28) were caught using the Suna trap (Figure 2A). In contrast to the *Anopheles* catch data, more culicines were caught in Mwanza compared to Lengwe (RR = 1.43, P = 0.041). When culicine data were pooled across villages there was no evidence for differences in the relative capture efficacy of the Suna trap (R = 0.76, P = 0.197) or HDT (R = 0.67, P = 0.060) against the HLC, however, performance was clearly dependent on the village of collection (Figure 2B).

**Figure 1.**
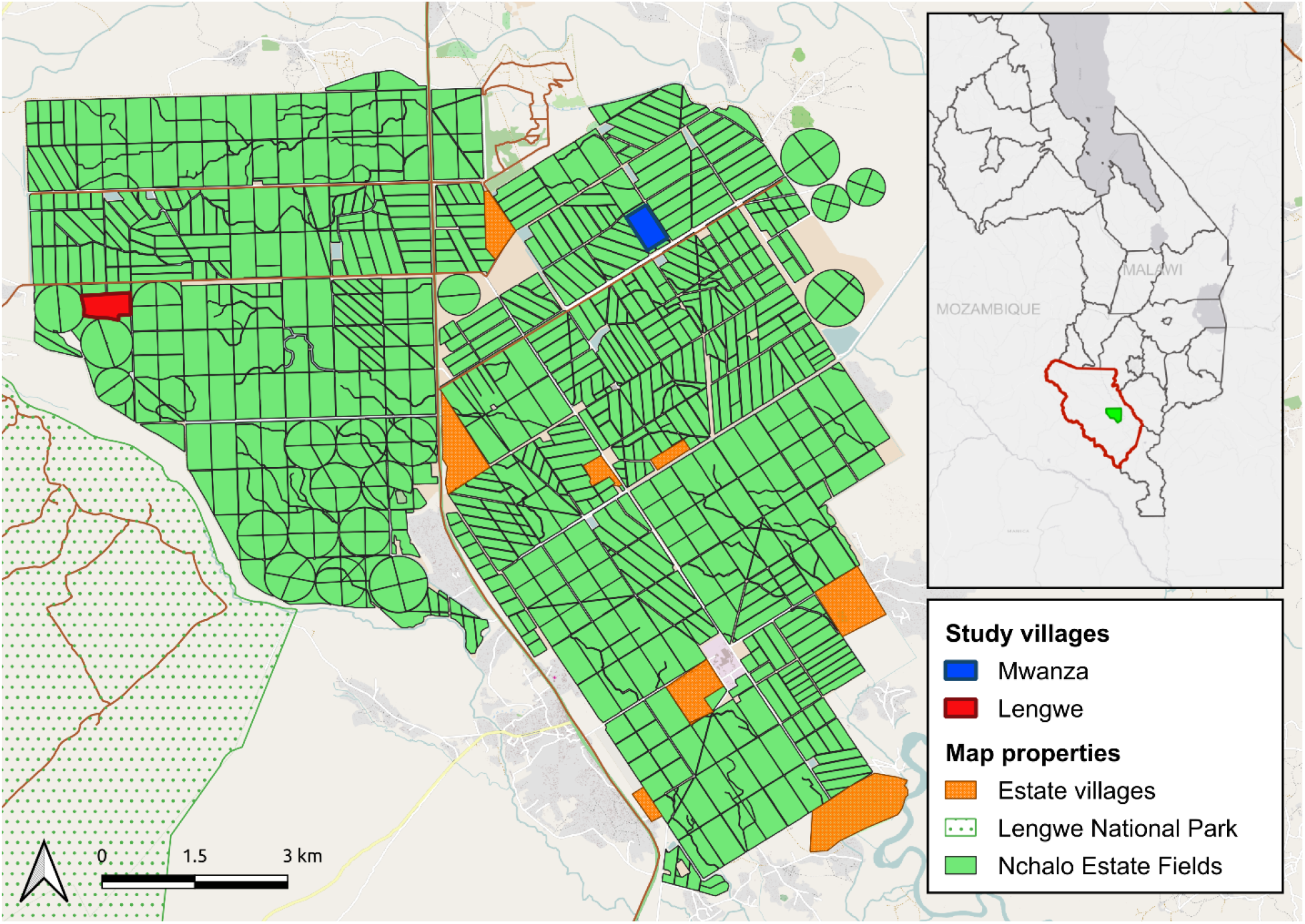
The location of the Illovo sugar estate within Malawi (inset) and study villages Mwanza and Lengwe.

**Figure 2.**
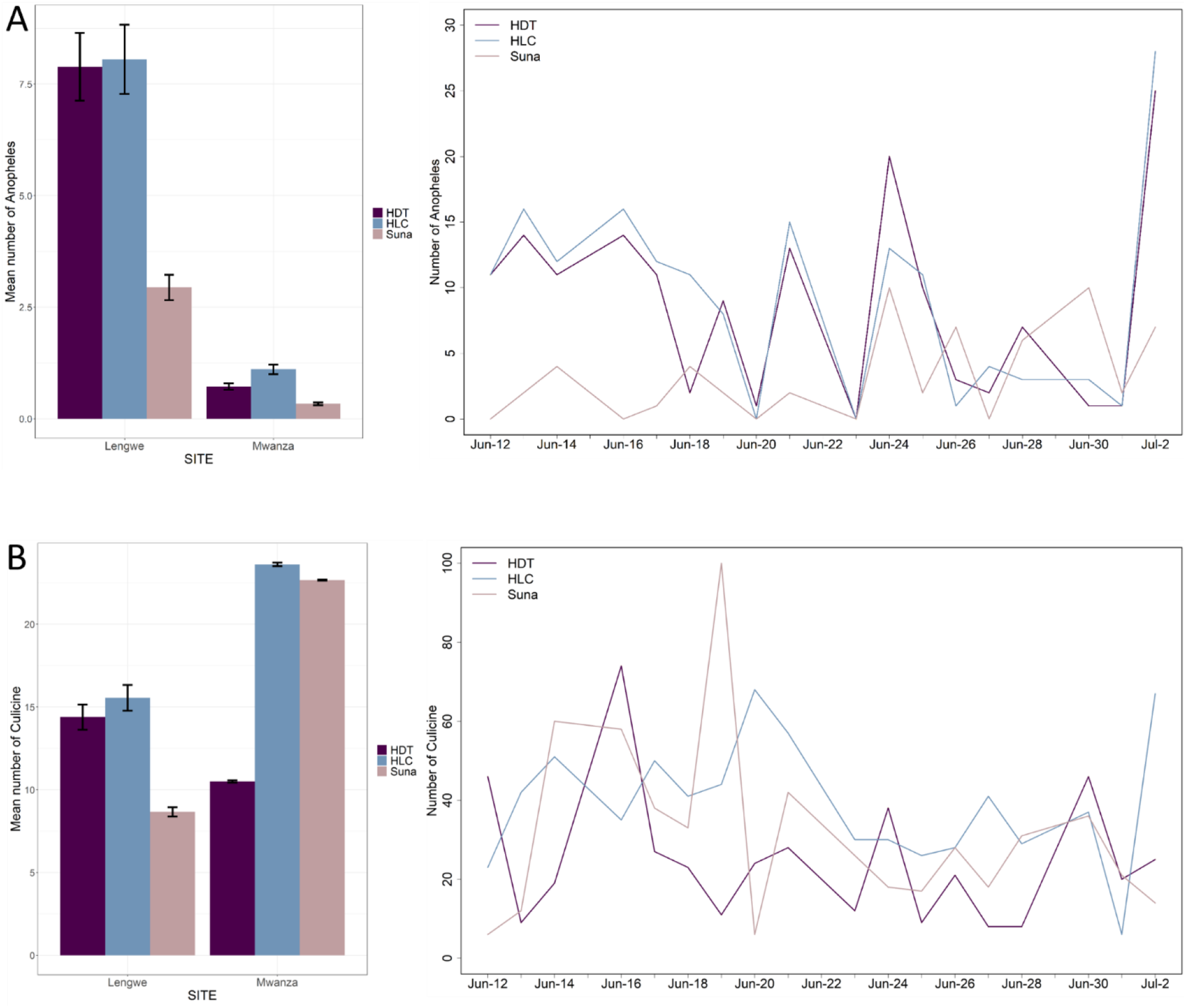
The mean number of mosquitoes caught per trap per night for each trapping method in Mwanza and Lengwe (± s.e.m) and a breakdown of the total number of mosquitoes caught per night in both villages. Data are presented for *Anopheles* (panel A) and culicines (panel B).

At the species level most *Anopheles* specimens were identified as *An. gambiae s*.*l*. (87%, n = 330) followed by *An. coustani* (11%, n = 40) and *An. funestus* (2%, n = 9). All *An. gambiae s*.*l*. underwent PCR to determine sibling species although we were unable to clarify 43 individual specimens due to non-amplification of PCR products. The proportion of unidentified *An. Gambiae s*.*l*. specimens caught per trap type was similar (HDT = 13.9%, HLC = 9.0%, Suna = 10.9%). Out of the remaining 287 *An. gambiae s*.*l*. samples, the predominant species was *An. arabiensis* (94.4%, n = 271) followed by *An. quadriannulatus* (4.2%, n = 12), with very few *An. gambiae s*.*s*. (1.4%, n = 4).

The relative proportions of *Anopheles* species did not substantially vary by trap and were consistent across Lengwe and Mwanza. *An. arabiensis* was the predominant species in all trap types in both villages although the relative proportions were slightly higher in the HDT (Lengwe = 0.74; Mwanza = 0.77) and Suna trap (Lengwe = 0.77; Mwanza = 0.83) compared with the HLC (Lengwe = 0.67; Mwanza = 0.65). This reflects the greater diversity of species caught using the HLC (Figure 3). In Lengwe, where *Anopheles* numbers were greater, the secondary vectors, *An. coustani* and *An. tenebrosus*, comprised 0.10 and 0.08 of HLC collections respectively; relatively higher than the proportion caught by either the HDT (0.03 and 0.01) or Suna trap (0.04 and 0.04). The very low numbers of other anopheline vectors caught in this study (*An. gambiae s*.*s*., *An. funestus* and *An. quadriannulatus*) make comparisons for these species trivial, although it is noteworthy that only the HDT and HLC caught *An. quadriannulatus*.

**Figure 3.**
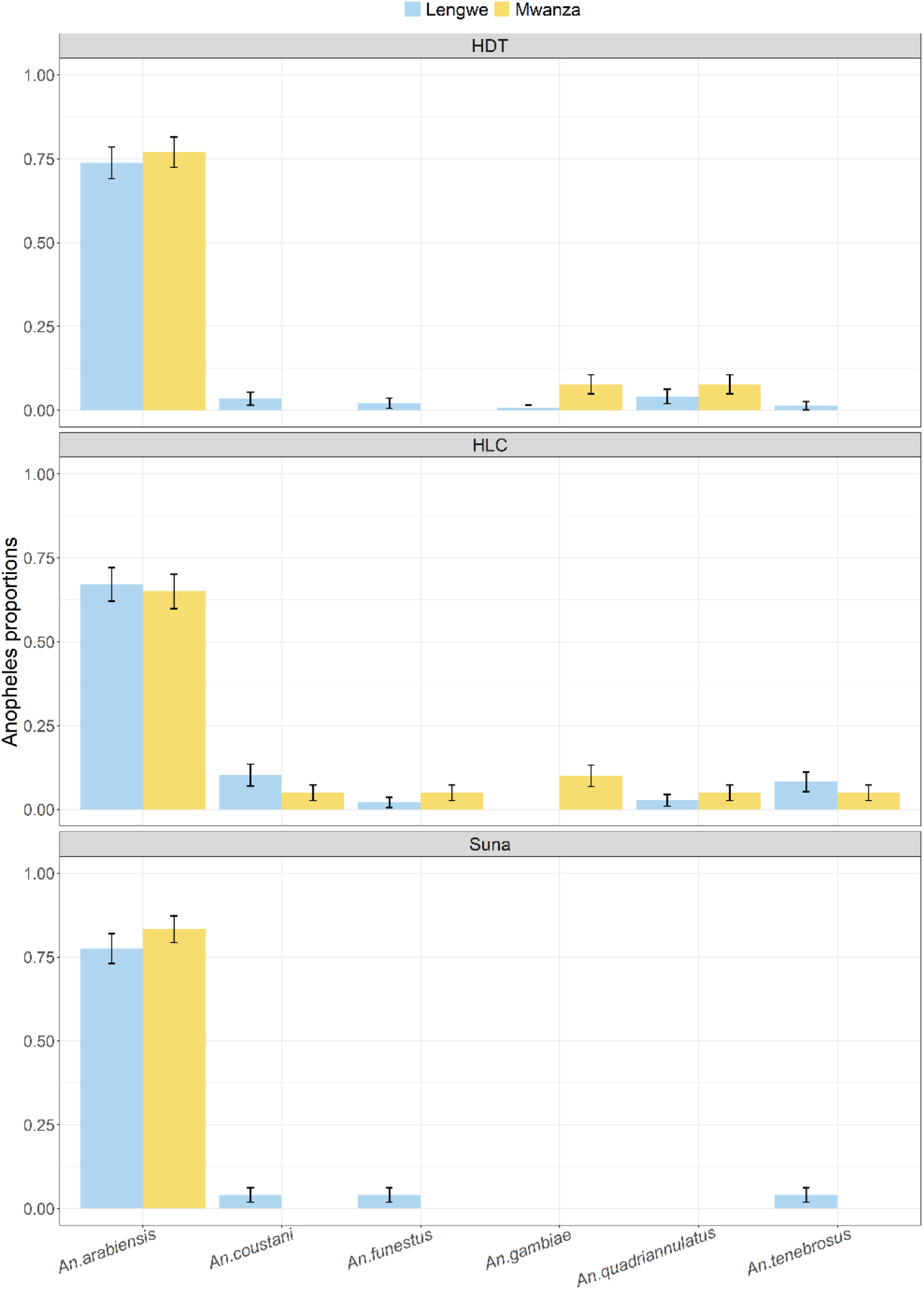
Relative *Anopheles* species composition (proportions ± s.e.m) for the HDT, HLC and Suna trap in the Nchalo sugar estate, Malawi. Data are presented for each study village.

All the *Anopheles* mosquitoes collected were females and the abdominal status were classified as either fed, gravid or unfed, although only six gravid mosquitoes were caught in the entire study. Over one-fifth of female *Anopheles* caught by HLC had taken a feed prior to collection (21.4%), much higher than the HDT (9.6%) and Suna trap (3.6%).

We investigated whether the sampling efficiency of each trap varied with *Anopheles* density using a modified test for linearity^19–21^. The density dependence (ß) and the sampling efficiency (α) parameters for the linear and non-linear models are provided in Table 1.

**Table 1.** Summary of the model parameter estimates for each trapping comparison.

| Model comparison | Location | Model 1 | Model 2 |  |
| --- | --- | --- | --- | --- |
| HDT vs Suna | | $\alpha_s$(95% CI) | $\alpha_s$(95% CI) | $\beta_s$(95% CI) |
|  | Lengwe | 0.38 (0.26, 0.50) | 8.56 (0.38, 13.40) | 0.27 (0.12,0.98) |
|  | Mwanza | 0.43 (0.15,1.17) | 0.44 (0.00, 430) | 0.05 (0.00,0.87) |
| HDT vs HLC | Lengwe | 1.03 (0.80, 1.30) | 0.65 (0.34,1.68) | 1.23 (0.83,1.82) |
|  | Mwanza | 1.52 (0.83, 3.17) | 1.76 (0.67,14.41) | 0.88 (0.32,195) |
| Suna vs HLC | Lengwe | 2.75 (2.03, 3.73) | 0.41 (0.29,0.64) | 8.84 (3.01,31.47) |
|  | Mwanza | 3.38 (1.50, 9.30) | 2.88 (1.01,41.63) | 1.25 (0.21,10.48) |

Figure 4 shows the fitted linear (model 1) and non-linear (model 2) relative sampling efficiencies for each trap comparison. When considering the density effects in model 2, the only curve which remains close to a straight line is for the HLC and HDT (Figure 4b). The mean catch per night for the HLC and HDT was closely correlated (Figure 2a) and for both Lengwe and Mwanza the 95% credible intervals for ß include the value for density independence (ß = 1) (Table 1). We note, however, that the fewer number of mosquitoes caught in Mwanza (n = 36) lead to extremely wide credibility intervals (particularly in the power model) and demonstrate the limit of this approach when mosquito catches are so few in individual sites. Nevertheless, across the two study villages, the fitted curves and density dependence estimates show a broad agreement between the sampling efficiencies of the HDT and HLC compared with the relationship between these two methods and the Suna trap.

**Figure 4.**
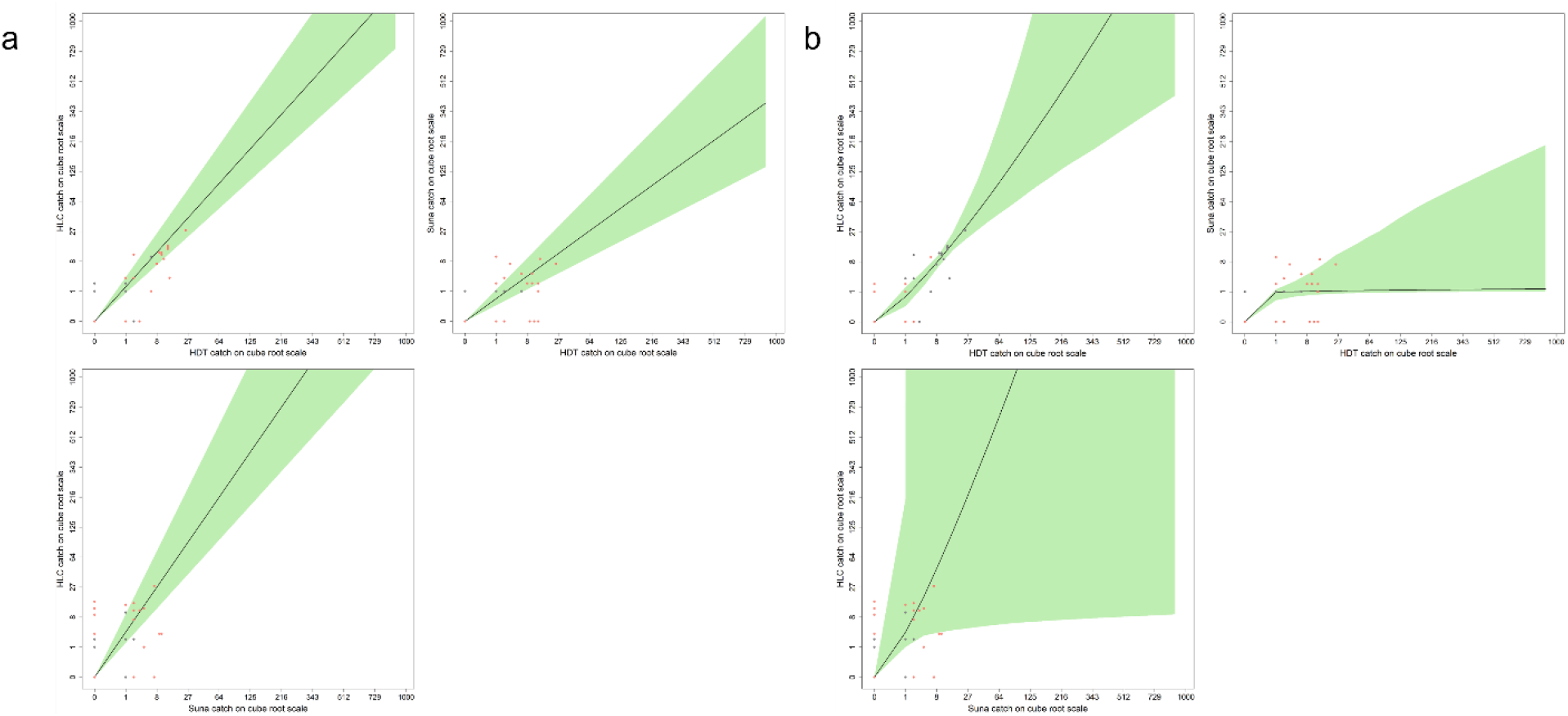
The fitted relative sampling efficiencies for each trap comparison. For each panel the number of mosquitoes caught per night are plotted against each other for the HDT vs. HLC (top left), HDT vs. Suna (top right) and HLC vs. Suna (bottom left). Sampling efficiencies are shown for the linear model in panel a and for the non-linear (power) model in panel b. The shaded green area corresponds to 95% credible intervals for the best fitting curve for each model comparison.

## Discussion

The ability to reliably monitor mosquito host-seeking behavior is an essential component of evaluating vector control. While most infectious bites continue to occur indoors^22^, quantifying the level of outdoor transmission has become increasingly important, particularly in parts of East Africa where there is now extensive evidence for shifts in biting behaviour^11^ and where the highly adaptable *An. arabiensis* has become the dominant vector^23^. Trapping methods are required that capture representative samples of the outdoor host-seeking *Anopheles* population, but entomological monitoring remains highly reliant on the gold-standard but ethically flawed HLC^17^. In this study, we add further evidence that the human baited HDT can be an efficient method for catching outdoor *Anopheles* vectors. We discuss the results in the context of sampling mosquitoes during the dry season in the lower Shire Valley of southern Malawi and within an extensive irrigated landscape.

Based on *Anopheles* catches per night, and within both villages on the Nchalo sugar estate, the HDT caught similar numbers of *Anopheles* as the HLC and outperformed the Suna trap (Figure 2). The predominance of *An. arabiensis* as the main vector (82% of identified anophelines and 94% of all *An. gambiae* s.l.) caught during our study means that few species-specific conclusions can be made concerning the diversity of vectors caught by each trap, although we note that the HLC did catch small numbers of a greater variety of anophelines including *An. gambiae s*.*s*., *An. coustani* and *An. tenebrosus*.

The comparative evaluation of mosquito trapping methods is context specific and depends on the diversity of local vector species, the environment and season. Rarely, if ever, will mosquito traps demonstrate an equivalent efficiency across all environmental contexts, and this is certainly the case for the HDT. In Burkina Faso, the HDT consistently outperformed the HLC regardless of season or mosquito genera with HDT catches of the main anthropophilic and endophagic vector, *An. coluzzi*, ten-fold higher than with the HLC^16^. In western Kenya, however, where *An. arabiensis* and *An. gambiae s*.*s*. are the dominant vector species, trap catches were dependent on the type of bait used^24^ with cattle-baited HDT catching 7-fold more *Anopheles* (mainly *An. arabiensis*) than the HLC. In contrast, when the HDT was baited with human odour, the HLC caught approximately 6-fold more *Anopheles*. The efficacy of the HDT may therefore be linked to species-specific differences in behavior; a conclusion supported by collections in the island of Sulawesi, Indonesia where a greater diversity of *Anopheles* species exists^25^.

The HLC and HDT caught between 2- and 4-fold more *Anopheles* than the Suna trap depending on the village. The Suna trap has shown promise as a relatively inexpensive device for both indoor and outdoor mosquito collections in laboratory and semi-field trials^26^. In recent field evaluations as part of the Majete Malaria Project - located approximately 30 kilometers away from Nchalo - the sampling efficiency of the Suna trap was similar to the HLC for both indoor and outdoor anophelines^18^. We note, however, that the relative proportion of the vectors in Majete (ratio of *An. gambiae s*.*l*.: *An. funestus* = 60:40) differs to those from Nchalo, providing a possible explanation for the discrepancy in trap performance.

We caught 4.5-fold more culicines than anophelines over the course of the 18 days of the study. The capture of relatively greater numbers of outdoor culicines compared to *Anopheles* is not unusual in field evaluations of mosquito traps. Indeed, similar findings were observed with the HLC and Suna trap in nearby Majete^18^. In contrast to the anopheline data, the overall pattern of culicines caught by each trapping method contrasted between the two villages (Figure 2). As we did not distinguish culicine specimens to species level we cannot make any robust conclusions about the relative efficacy of each trap to sample culicines, but we hypothesize that the differential catches in Lengwe and Mwanza are due to the species composition.

The HLC requires volunteers to manually aspirate mosquitoes as they land to take a blood-feed. This exposes volunteers to potentially infectious mosquito bites which presents ethical concerns^17^. Aside from the safety issues, the data collected by HLC volunteers varies by individual, imposing a layer of experimental bias^27^. Alternatives to the HLC must be able to collect enough mosquitoes to justify their use and demonstrate the same functionality in terms of indoor/outdoor biting, have stable capture rates across the night and comparable efficiency irrespective of local mosquito densities^28,29^. Due to the nature of our study design, we did not address all these criteria, but we were able to investigate density dependence. Two traps are said to be density dependent if the relative sampling sensitivity varies with mosquito density. For example, in the case of the mosquito electrocuting trap (MET), which shows promise as an outdoor sampling tool, the relative sensitivity of the MET and HLC to catch *Anopheles* is largely unaffected by mosquito density and the traps can be considered density independent^19,20^. Here, we show that the HDT and HLC have broadly comparable sensitivities for catching an outdoor vector population consisting largely of *An. arabiensis*, irrespective of density.

Irrigated sugar fields are a perennial source of habitat for anopheline vectors. In cultivated sugar farms across eastern and southern Africa, *An. arabiensis* is one of the main beneficiary species, thriving in the transient and dynamic surface waters provided by irrigation channels and drainage systems. Plenty of reports exist documenting higher mosquito densities and infection rates in localities in proximity to irrigated agricultural land dominated by *An. arabiensis* from Kenya, Ethiopia and Tanzania ^6,7,9^. The finding that *An. arabiensis* is the main outdoor host-seeking vector within Nchalo during the dry season is significant as this will impact current vector control efforts. Since 1990, Illovo has conducted a private IRS programme to protect its employees and their families, and now routinely conducts an annual spray of all houses within the estate with a long-lasting application of pirimiphos-methyl (Actellic 300CS). An analysis of monthly malaria health facility data over a five-year period shows that IRS is having a positive impact on reducing malaria cases, but it is almost certain that residual outdoor transmission is occurring and is most likely driven by *An. arabiensis*. Year-round monitoring of indoor and outdoor mosquito behaviour in the area is needed to determine how to effectively tailor vector control efforts in such a landscape.

We propose that the HDT is an efficient method for sampling outdoor host-seeking anophelines during the dry-season in southern Malawi. Further investigations are needed to make comparisons with the HLC in other environmental contexts within Malawi to ensure that samples caught by the HDT are representative of local populations and catches remain stable throughout the night. While this study was conducted during just a single dry season, the predominance of *An. arabiensis* suggests that the irrigated sugar farms provide a year-round refuge for this species and efforts should be directed towards understanding the behavior of *An. arabiensis* particularly in the context of the local IRS programme.

## Methods

### Study site

Illovo Sugar is the major producer of sugarcane in the Southern African Development Community (SADC) region. In Malawi, Illovo has two cane growing sites producing 2 million tonnes of sugarcane combined per year. Our study was conducted in two villages, Mwanza (−16.1995; 34.8009) and Lengwe (−16.1869; 34.8805), within the Nchalo estate which sits within the low-lying Shire Valley of the southern district of Chikwawa (70 m.a.s.l.) (Figure 1). Temperatures in the valley can rise up to 40°C during the dry season which lasts from May to November while peak rainfall occurs sometime between December and February^30^. The two major malaria vectors found in the regions are *An. arabiensis* and *An. funestus*, their relative proportions varying seasonally^31^. The Nchalo estate incorporates ∼13,000 ha of sugarcane plots irrigated by a range of methods supplied by water from the Shire River. Illovo has conducted an IRS programme since 1990 using a variety of pyrethroid-based compounds until 2014 before a switch to a single annual application of Actellic 300 CS (pirimiphos-methyl) prior to the onset of the rainy season^15^. Households within Nchalo receive bednets as part of the National Malaria Control Programme distribution campaigns.

### Study design and mosquito sample collection

We selected three houses within each of Mwanza and Lengwe for mosquito sampling. All houses were roofed with iron sheets and had fully closed eaves. Sampling houses were approximately 100 meters apart to reduce the influence of nearby traps on mosquito collections at other houses. Mosquito sampling was performed using three different methods: the HDT, HLC and Suna trap. All mosquito collections were conducted in the outdoor environment and traps were set-up approximately three metres within the compound of selected houses. For our study we used a repeated 3 × 3 Latin square experimental design (trap x house x night). Two concurrent Latin square rotations were conducted within each village (i.e., six houses containing either HDT, Suna trap or HLC during any single night). The two Latin squares were repeated six nights a week for three weeks (18 nights of data collection per village and 36 trap nights) between the months of June and July 2019. All experiments began at 1800 and ran until 0600 the following morning.

A standardized version of the HDT (developed by the University of Greenwich and Biogents) with some minor modifications was used in all experiments. In brief, the HDT uses a protected host as bait from which odour is funnelled down a 6 m PVC pipe towards a visually contrasting black and thermally heated trap. A transparent adhesive plastic sheet (Barrettine Environmental Health, Bristol, UK) covers the circumference of the trap to catch mosquitoes as they land. During each collection night a volunteer slept in a tent, positioned approximately five meters from the house, as a source of odour. At the end of the collection period, the adhesive sheets were transported to the laboratory and mosquitoes removed using forceps and non-toxic Mobe-Moat solvent (Barrettine Environmental Health, Bristol, UK). Suna traps were suspended outside the house approximately 30 cm above ground level^26^. Suna traps were baited with the MB5 attractant blend, suspended outside the house approximately 30 cm above ground level and supplied with CO2 produced from yeast and molasses (from the Illovo estate) for fermentation. Mosquitoes were collected from the HDT and Suna traps between 0700 and 0900.

Four HLC collectors were recruited from Mwanza and Lengwe and trained prior to the study. All collectors were given malaria prophylaxis (doxycycline) at a daily dose of 100mg for the duration of collections and for five days before and after the study. Two volunteers were assigned per house with each working one shift per night (1800-0000, 0000-0600) and taking a 15-minute break per hour. Collectors were sitting approximately 3 m from the house with their legs exposed from their knees down. Mosquitoes landing on their legs were collected using a mouth aspirator. To minimize individual host-attraction bias, after six hours (mid-point) during a single collection night, HLC and HDT volunteers swapped places.

Mosquitoes were stored individually, separated by sex and morphologically identified to genus level^32^ to differentiate anopheline and culicine catches. All anophelines were identified by species complex and sub-species level using PCR^33^ and classified as either fed or unfed. Ethical approval for this study was obtained from the College of Medicine Research Ethics Committee (COM-REC) (No. 2605).

### Data analysis

The primary outcome was the mean number of mosquitoes collected per trap per night. Data was analyzed in R version 3.5.1^34^. Generalized Linear Mixed Models (GLMM) were used to compare the mean number of mosquitoes collected per trap per night with a negative binomial distribution to account for over-dispersion. Models were formally compared using Akaike Information Criteria (AIC) and log-likelihood ratio tests (LRT). Trap type and village were considered independent fixed effects with the night of collection fit as a random effect. Separate models were fit for nightly collections of *Anopheles* and Culicine mosquitoes.

The number of mosquitoes caught by two trapping methods targeting the same population should be roughly proportional across the range of mosquito densities. To estimate the comparative sampling efficiencies of each trap we followed the approach by Briet *et al*.^21^ and fitted the linear model below:

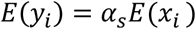

In this model, *E*(*y*_*i*_) is the expected number of mosquitoes caught by a trap Y during the night; *E*(*x*_*i*_) is the expected number of mosquitoes caught by trap X on the same night; *α*_*s*_ is the relative sampling efficacy corresponding to site *s* (Lengwe or Mwanza). It is assumed that the mosquito counts in each trap follows a Poisson distribution such that *x*_*i*_ *∼ Poisson*(*E*(*X*_*i*_)) and *y*_*i*_ *∼ Poisson*(*E*(*Y*_*i*_)). Mosquito densities were assumed to follow a log-normal distribution, i.e. *ln*(*E*(*X*_*i*_))*∼N*(*μ, σ*^2^). Three models were fitted for each pair of traps (i.e. HDT with HLC, HDT with Suna and HLC with Suna) and only *Anopheles* data were assessed.

For the density dependent analysis, the linear model was modified as follows:

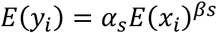

The parameter *β*_*s*_ is the site-specific exponent governing the relationship between the two traps. Density independence (or proportionality) is achieved when *β*_*s*_ approximates 1 and density-dependent effects are considered if *β*_*s*_ is different from 1.

Model fitting was done in a Bayesian framework using Markov chain Monte Carlo algorithm in WinBUGS 1.4 through the R2WinBUGS R interface. We assigned non-informative uniform distribution priors to all the model parameters and constrained them to be positive. For each model, we ran 40,000 iterations, a burnin of 20,000 and a thinning parameter equal to 20 thus giving 1000 posterior samples for inference.

### Ethical approval

The study was conducted with ethical approval from the College of Medicine Research and Ethics Committee (COMREC) (P.02/19/2605) in Malawi. All study procedures were performed in accordance with the relevant national guidelines. Prior to the commencement of the study, we obtained support from the District Health Officer of Chikwawa and from village leaders in Mwanza and Lengwe. Written informed consent was obtained for all volunteers and householders recruited for trap operations. Study information including the purposes, benefits and risks was provided to all participants in both English and Chichewa.

## Acknowledgements

We thank the volunteers and households who participated in the mosquito sampling in Lengwe and Mwanza. We are grateful to Dr Albert Mkumbwa and Illovo Sugar Ltd for permission to conduct research on the Nchalo estate. KZ was supported by an institutional training grant awarded as part of the Wellcome Strategic award number 206545/Z/17/Z to the Malawi-Liverpool-Wellcome Trust Clinical Research Programme (MLW), administered under the joint MLW/Kamuzu University of Health Sciences Training Committee.

## Author contributions

KZ and CJ conceived the study; KZ, CJ and FMH designed the study; KZ, LD and PN performed field and lab experiments; JC, KZ, DM and CJ analysed the data; KZ and CJ wrote the manuscript; all authors read, commented on, and approved the final manuscript.

## Additional information

The authors declare no competing interests.

